# Substrate accessibility of bovine milk proteins to a *Pseudomonas* metalloprotease reveals why caseins are more readily hydrolysed than whey proteins

**DOI:** 10.64898/2026.10.02.756230

**Authors:** Aritra Sinha, Tiziana Racca

## Abstract

Structure-aware proteolysis prediction often ranks candidate cleavage sites using solvent accessibility from static protein structures, yet this assumption is rarely tested against native-state cleavage data. We examined this problem using AprX, a serralysin-family protease that readily hydrolyses caseins but shows lower activity toward native whey proteins. Across six bovine milk proteins, sequence compatibility and local scissile-bond accessibility did not separate caseins from whey proteins. In contrast, accessibility fell progressively in whey proteins as the required contiguous substrate window increased, reaching 0% across the working P4-P4′ footprint, while all evaluable casein windows remained accessible in isolated-chain ensembles. Local remodelling showed that whey segments could still adopt AprX-compatible backbone geometry. An external thermolysin benchmark rejected a rigid whole-protein clash metric, but did not alter the contiguous-accessibility result. These findings identify footprint-scale accessibility as a distinct structural variable and show why single-residue solvent exposure can overestimate protease accessibility in folded proteins.

## 1. Introduction

Psychrotrophic *Pseudomonas* spp. is a persistent concern in refrigerated raw milk because they can grow during cold storage and secrete extracellular proteases before thermal processing. Although the cells are readily inactivated by pasteurisation or ultra-high-temperature (UHT) treatment, some extracellular proteases are considerably more heat resistant. AprX, an extracellular alkaline zinc metalloprotease of the serralysin family, is a prominent example and has been repeatedly associated with proteolysis, aggregation, gelation and sedimentation leading to shelf-life failures and premature spoilage in UHT milk (Marchand et al., 2009; Baglinière et al., 2013; Zhang et al., 2018; Zhang et al., 2020). Purified AprX has been shown to hydrolyse all four major caseins and destabilise UHT-treated casein micelles, while κ-casein hydrolysis has been associated with gel formation during storage (Baglinière et al., 2013; Zhang et al., 2018). AprX-mediated proteolysis therefore remains an important mechanism of UHT milk spoilage, although aspects of its biochemical action remain unresolved (Zhang et al., 2019; Almeida et al., 2026).

Milk proteins are not equally susceptible to AprX. Matéos et al. (2015) reported extensive hydrolysis of caseins but not whey proteins by AprX from *Pseudomonas* LBSA1. Numerous cleavage sites were identified in α_S1-_, β-and κ-caseins, indicating broad specificity, although positional preferences were evident, with Lys, Arg and Phe favoured at P1 and Asp, Glu and Pro less favourable. Subsequent peptidomics similarly identified preferential cleavage around His, Lys, Arg, Leu, Met and Phe at P1/P1′, whereas Asp, Glu and Cys at either position, Ile at P1 and Pro at P1′ were comparatively unfavourable (Zhang et al., 2023). Overall, it can be stated that AprX preferentially accommodates hydrophobic and aromatic residues at P1/P1′, while also favouring certain basic side chains. However, β-lactoglobulin-derived peptides have also been detected following AprX treatment under other conditions, indicating that whey-protein resistance is relative and dependent on structural and processing context and can’t be explained solely from absolute sequence incompatibility (Zhang et al., 2023).

This difference has commonly been attributed to the difference between the protein conformation. Caseins are intrinsically disordered proteins that populate dynamic ensembles, although in milk they associate within organised casein micelles (Holt et al., 2013). In contrast, β-lactoglobulin is a compact β-barrel lipocalin and α-lactalbumin is a globular Ca²⁺-binding protein stabilised by tertiary interactions and disulfide bonds (Kontopidis et al., 2004; Permyakov et al., 2016). Despite this, globular folding, however, does not rule out possibility of proteolysis. Studies have shown that folded proteins can undergo limited cleavage at exposed and flexible regions when local backbone rearrangement permits access to the protease active site (Hubbard et al., 1994; Fontana et al., 2004), as demonstrated by selective proteinase K cleavage of an exposed loop in native avidin without global unfolding (Ellison et al., 1995). Thus, which structural property of the fold prevents otherwise compatible regions from being productively presented to AprX remains unanswered.

Large-scale analyses have identified the residues surrounding the scissile bond, solvent accessibility, flexibility, secondary structure and local unfolding propensity as distinct determinants of cleavage (Kazanov et al., 2011; Belushkin et al., 2014). Moreover, proteases generally recognise substrates in extended strand-like conformations across several subsites rather than only at the two residues flanking the scissile bond (Hubbard et al., 1994; Madala et al., 2010). Consequently, a bond may be solvent exposed while the neighbouring backbone remains unable to adopt the extended geometry required for productive binding.

This principle is relevant to serralysins, the class to which AprX belongs. Louis et al. (1998) showed that *Pseudomonas aeruginosa* serralysin tolerated multiple residues at P1 and P1′ but hydrolysed longer peptide substrates much more efficiently, supporting recognition across an extended series of subsites. Such substrate-side effects can be important even when the protease itself is unchanged. For example, native β-lactoglobulin is comparatively resistant to pepsin and trypsin, whereas heating, denaturants or reduction markedly increase its susceptibility by exposing previously inaccessible regions (Guo et al., 1995). Processing, aggregation, mineral interactions or other changes in milk-protein organisation could therefore alter AprX susceptibility independently of changes in AprX abundance or catalytic competence.

Despite this, previous studies have not distinguished whether resistance to AprX arises primarily from unfavourable sequence chemistry, burial of the scissile bond, failure of an extended substrate segment to become simultaneously accessible, or steric exclusion of the intact globular protein from the catalytic cleft. Existing work has focused predominantly on AprX occurrence, regulation, heat stability, detection, activity and cleavage specificity (Zhang et al., 2019; Almeida et al., 2026), leaving the structural contribution of the substrate less well defined.

Accordingly, this study tested whether the differential susceptibility of bovine caseins and whey proteins to AprX can be explained by sequence compatibility, local cleavage-site exposure or accessibility of an extended substrate segment. Structural modelling was used to define a working AprX substrate footprint, which was evaluated across the major bovine milk proteins using sequence scoring, structural-accessibility analysis and structure-guided peptide modelling. Finally, local conformational-accommodation modelling was used to test whether exposed whey regions could adopt the AprX-bound geometry while remaining attached to the parent fold. A whole-protein steric descriptor was evaluated as an exploratory extension and was subsequently challenged using an external thermolysin benchmark with independently known native-state cleavage outcomes.

## 2. Materials and methods

### 2.1. Study design and overall workflow

A computational workflow was developed to investigate whether the differential susceptibility of bovine caseins and whey proteins to the heat-stable metalloprotease AprX could be due to differences in peptide-sequence compatibility, structural accessibility, or presentation of an extended substrate segment to the enzyme active site. The analysis combined an AprX structural model, experimentally determined structures of homologous serralysin-family metalloproteases, experimentally determined bovine whey-protein structures, conformational ensembles of intrinsically disordered caseins, experimentally identified AprX cleavage sites, residue-level solvent-accessibility calculations, and structure-guided peptide modelling.

Six major bovine milk proteins were analysed: α_S1-_casein, α_S2-_casein, β-casein, κ-casein, β-lactoglobulin and α-lactalbumin. Every peptide bond within the mature sequences of these proteins was considered as a potential cleavage position, resulting in 1,063 candidate peptide bonds. Analyses were performed sequentially to determine whether sequence compatibility or structural accessibility could account for the observed difference in susceptibility between caseins and whey proteins. The overall analytical workflow is summarised in Figure 1.

**Figure 1.**
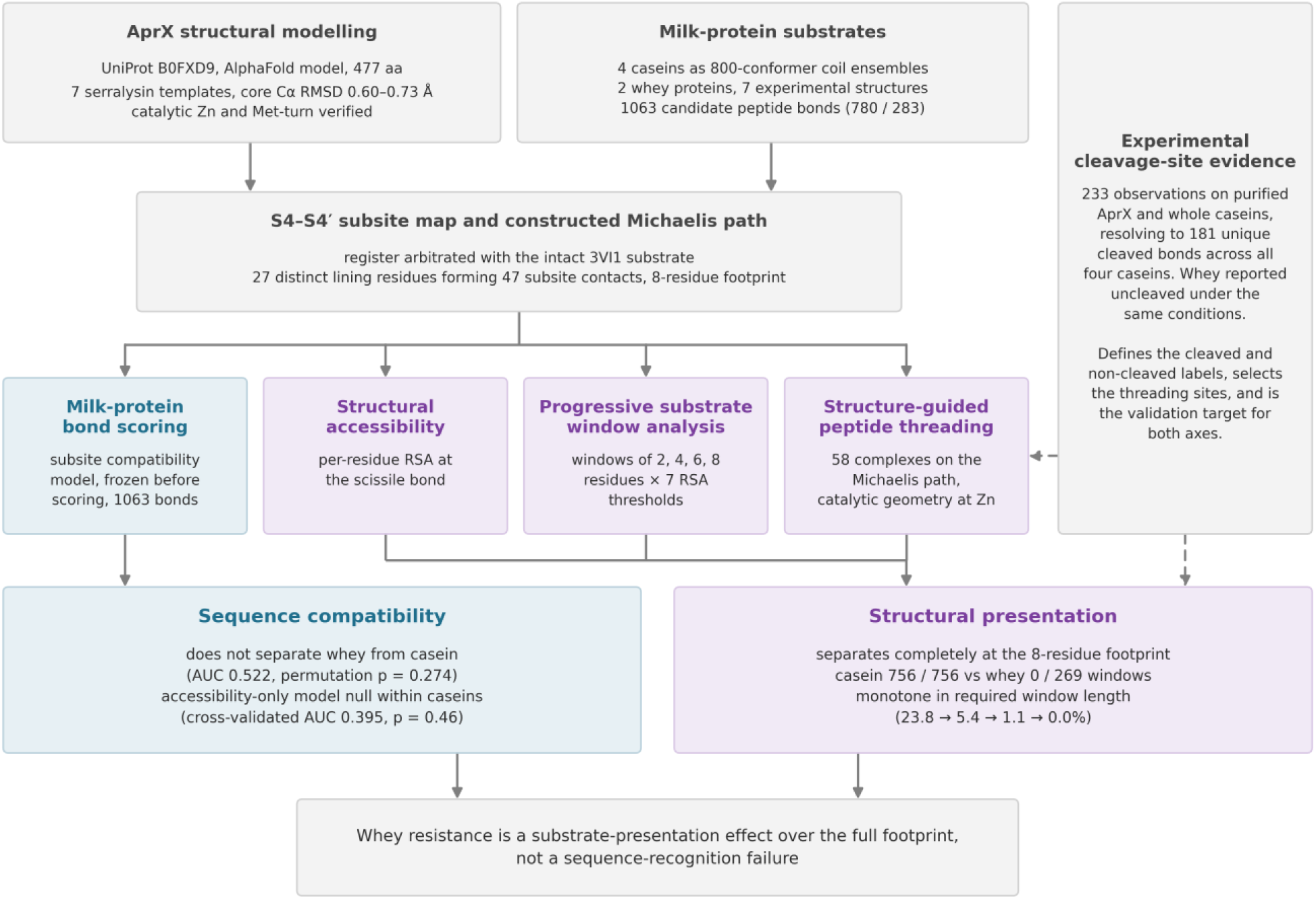
Overview of the study workflow. AprX structural modelling was combined with milk-protein bond scoring, structural accessibility analysis, progressively longer substrate-window analysis, structure-guided peptide threading and experimental cleavage-site evidence to distinguish sequence compatibility from structural presentation of the substrate.

**Figure 2.**
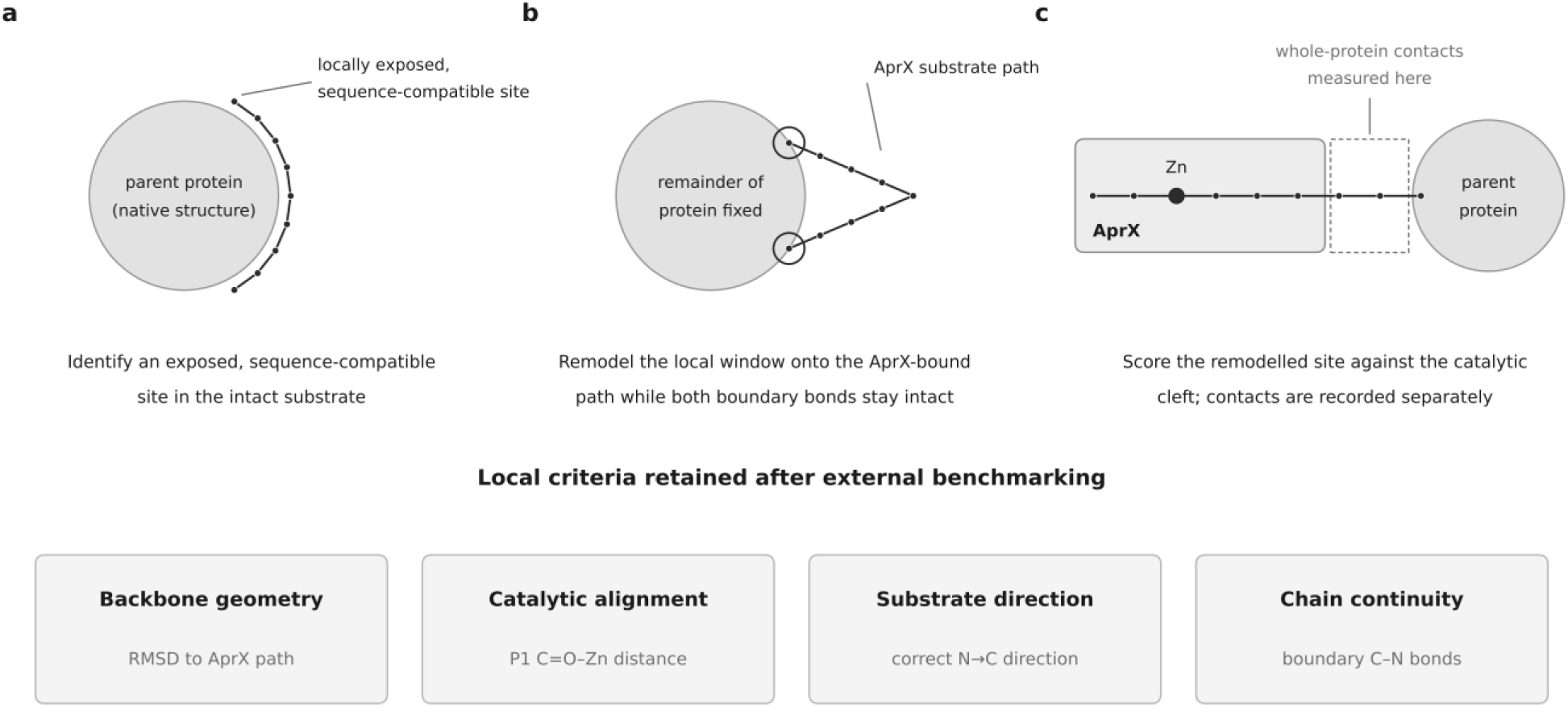
Schematic overview of the conformational-accommodation analysis and subsequent benchmarking. (a) Sequence-compatible, locally exposed regions were identified within the native parent protein. (b) Each candidate segment was remodelled onto the AprX substrate path while preserving covalent continuity with the parent fold. (c) Local backbone geometry, catalytic Zn²⁺ alignment, substrate direction and chain continuity were evaluated after remodelling. Whole-protein contact burden was recorded separately and externally benchmarked rather than retained as an absolute pass/fail criterion.

### 2.2. AprX sequence and structural model

The AprX sequence from *Pseudomonas fluorescens* (UniProt B0FXD9) was used as the enzyme reference. The AlphaFold Database model (version 6) was used as the working structure, as no experimental structure was available for this protein, (Jumper et al., 2021; Varadi et al., 2022). Before use, the model was compared with seven experimentally determined serralysin-family structures (PDB 1KAP, 1AKL, 1SMP, 1SAT, 1AF0, 1JIW and 1SRP). Structures were aligned using least-squares Cα superposition based on the Kabsch algorithm (Kabsch, 1976), and agreement was assessed from Cα RMSD, sequence identity and conservation of the catalytic and Zn²⁺-binding residues. The position of the catalytic Zn²⁺ in the experimental structures was also compared after alignment. The AprX catalytic region showed high AlphaFold confidence and close agreement with the experimental serralysin structures and was therefore used for subsequent structural analyses.

### 2.3. Definition of the AprX substrate-binding footprint

The substrate-binding trajectory within AprX was inferred using structurally characterised serralysin-family enzymes containing bound ligands. PDB 3VI1, an experimentally determined *Pseudomonas aeruginosa* alkaline metalloprotease structure complexed with the N-terminal Substance P fragment RPKPQQ, was used as the principal structural anchor (wwPDB: pdb_00003vi1).

The 3VI1 structure was aligned to the AprX model using the conserved catalytic core. Following superposition, the bound peptide was transferred into the AprX coordinate system to define the orientation of the substrate backbone relative to the catalytic zinc ion and surrounding binding subsites. The transferred complex aligned over 419 core Cα atoms with a structural RMSD of approximately 0.65 Å.

Substrate positions were assigned according to the Schechter and Berger nomenclature. The central substrate trajectory directly supported by the 3VI1 peptide extended approximately from P4 to P2’. Additional serralysin-family ligand structures were used to extend the working map through P3’ and P4’. The resulting P4 to P4’ interval was used as an eight-residue working structural footprint around the scissile bond.

### 2.4. Milk-protein sequences and candidate peptide bonds

Bovine protein sequences were obtained from UniProt (α_S1-_casein, P02662; α_S2-_casein, P02663; β-casein, P02666; κ-casein, P02668; β-lactoglobulin, P02754; and α-lactalbumin, P00711), and signal peptides were removed. Every peptide bond in the mature UniProt sequences was systematically designated as a potential AprX cleavage site, yielding 1,063 candidate bonds: 780 from the four caseins and 283 from β-lactoglobulin and α-lactalbumin. Terminal positions for which complete sequence windows could not be obtained were retained for analyses that did not require the full window and excluded only where the corresponding structural window was required. Experimentally reported AprX cleavage sites were mapped to the mature milk-protein sequences, and duplicate cleavage sites were removed before analysis.

### 2.5. Structure-derived sequence-compatibility scoring

For each P4–P4′ octapeptide, a custom structure-derived compatibility score was calculated for each candidate P4–P4′ substrate window to estimate how well its residues matched the corresponding subsites within the AprX catalytic cleft. The score combined five physicochemical features: steric fit between residue size and available subsite space, under-filling of the subsite, electrostatic complementarity, desolvation of charged residues and hydrophobic burial. Subsite capacity was estimated from the residue occupying the equivalent position in homologous ligand-bound structures together with the remaining local pocket volume. Residue side-chain volumes were taken from the Zamyatnin scale, hydrophobicities from the Fauchère–Pliska scale, and side-chain charges were calculated at pH 6.8 using Henderson–Hasselbalch relationships. The compatibility score was conceptually inspired by empirical molecular-recognition scoring functions such as AutoDock4, in which steric complementarity, electrostatic interactions and desolvation are combined to evaluate molecular compatibility (Huey et al., 2007; Morris et al., 2009). Here, these principles were adapted to residue–subsite matching within the AprX catalytic cleft, with an additional hydrophobic contribution.

Positional weights were designated to each subsite depending on the amount and quality of structural support available for each subsite. The central S2–S2′ positions were given full weight because they were directly occupied by the intact substrate in PDB 3VI1 and supported by additional serralysin ligand structures. S3′ and S4′ were given lower weights because their positions were inferred only from inhibitor-bound structures. S4 was not included in the compatibility score because it formed a solvent-exposed rim instead of a well-defined binding pocket. All term and positional weights were defined heuristically and were not fitted to experimental AprX cleavage data. Phosphoserine was included as a separate residue type, whereas phosphothreonine was represented as unmodified threonine because a corresponding parameter set was unavailable. Full equations, parameter values and weighting factors are provided in Supplementary Methods. The scoring model was subsequently evaluated against experimentally observed AprX specificity and was treated as a relative structural-compatibility index.

### 2.6. Structural preparation of whey proteins

Experimentally determined PDB structures were used to represent the folded whey proteins. β-Lactoglobulin was represented primarily by PDB 1BSQ, an X-ray structure of bovine β-lactoglobulin (Qin et al., 1999), with 4GNY, 7KOT and 1B8E used to test sensitivity to structural variation. α-Lactalbumin was represented primarily by PDB 9OZW, with 1F6S and 7EKA used as alternative structures. Protein preparation was done following standard structural-modelling procedures used in tools such as PDBFixer/OpenMM, including resolution of alternate conformations, removal of non-essential heteroatoms and crystallographic waters, and reconstruction of missing atoms within experimentally resolved residues (Eastman et al., 2017). Entire residues absent from the experimental structures were not modelled for the primary accessibility analysis, so that accessibility estimates were based only on experimentally resolved coordinates

### 2.7. Solvent-accessibility analysis of whey proteins

Solvent accessibility was calculated using the Shrake–Rupley method (Shrake and Rupley, 1973), as implemented in Biotite (Kunzmann and Hamacher, 2018), using a 1.4 Å solvent probe and ProtOr atomic radii. Absolute surface areas were converted to relative solvent accessibility (RSA) using residue-specific maximum values. Residue depth, secondary structure and local atomic packing were also calculated as complementary structural descriptors. For each candidate bond, local accessibility was defined from the RSA of the residues flanking the scissile bond.

### 2.8. Generation of casein conformational ensembles

Caseins were represented as multiple possible protein conformations because they are intrinsically disordered and do not adopt a single stable tertiary structure (Holt et al., 2013). Using statistical-coil approaches (Ozenne et al., 2012), 200 conformers were generated for each casein, giving 800 structures in total. These ensembles were intended to represent accessible isolated-chain conformations.

### 2.9. Comparable accessibility calculation for casein and whey proteins

To compare caseins and whey proteins on the same structural basis, both were represented using the same backbone and Cβ atoms before solvent-accessibility calculation. Solvent-accessible surface area was calculated using the Shrake–Rupley method (Shrake and Rupley, 1973) and converted to relative solvent accessibility by normalisation against the corresponding residue in a maximally exposed Gly–X–Gly reference peptide, following RSA approaches (Miller et al., 1987; Tien et al., 2013). For caseins, accessibility was calculated across all 200 conformers per protein and summarised across the ensemble; the same calculation was applied to the experimental whey-protein structures.

### 2.10. Contiguous substrate-window accessibility analysis

To test whether accessibility across a longer substrate-sized region separated the protein classes better than bond-level exposure, accessibility was examined over windows of 2, 4, 6 and 8 residues. This design was motivated by evidence that cleavage of folded proteins can require rearrangement of neighbouring residues (Hubbard et al., 1994), that proteases bind extended substrate segments across several subsites (Madala et al., 2010), and that serralysin hydrolysis increases with peptide length (Louis et al., 1998). The eight-residue window represented the working P4–P4′ AprX footprint. RSA thresholds from 0.05 to 0.35 were tested, with 0.25 used for primary reporting. This window analysis was developed after the initial bond-level result and was treated as a post-hoc mechanistic test.

### 2.11. Analysis of sequence-compatible and locally accessible whey sites

To test whether local exposure of the scissile bond was sufficient for AprX cleavage, 33 whey-protein sites that were both sequence-compatible and locally exposed were examined as counterexamples to a simple burial model. Accessibility at each site was compared across the full P4–P4′ window because of the earlier stated rationale that proteases generally require an extended substrate segment to enter the catalytic cleft rather than only an exposed bond (Hubbard et al., 1994; Madala et al., 2010). The lowest RSA within this window was used to identify neighbouring residues that could restrict presentation of the complete substrate segment.

### 2.12. Structure-guided peptide threading

Selected P4–P4′ peptides were removed from the intact whey-protein fold and modelled to test whether their sequences could adopt the AprX-bound geometry outside the parent fold. Their initial backbone geometry and orientation were taken from the experimentally observed substrate in PDB 3VI1. The complexes were then structurally relaxed. Geometry compatibility was assessed from scissile-bond positioning relative to catalytic Zn²⁺, peptide direction, steric fit and catalytic geometry.

### 2.13. Validation of the peptide-threading procedure and sensitivity analysis

The 3VI1 peptide was used as a positive control to confirm that the modelling workflow could reproduce the geometry on which it was based. Independent validation with the GSNS peptide from PDB 1KAP tested whether the same procedure could also reproduce a different experimentally observed metalloprotease-bound peptide, giving a backbone RMSD of ∼0.73 Å. Reversing the peptide orientation provided a negative control to confirm that incorrect binding direction was rejected. Because several poses lay just below the original Zn–O distance cutoff, 16 experimental metalloprotease structures were examined to determine whether the threshold was too strict; this supported 1.84 Å as a sensitivity limit, while 1.90 Å remained the primary criterion.

### 2.14. Conformational accommodation and external benchmark

To test whether local backbone constraints could explain whey resistance, nine sequence-compatible, locally exposed whey sites and eight experimentally cleaved casein sites were remodelled while remaining attached to their parent proteins, following the conceptual approach of Hubbard et al. (1994). Windows of 8, 12, 16 and 20 residues around each cleavage site were allowed to move in OpenMM using the Amber14 force field with GBn2 implicit solvent, while the remainder of the protein was fixed. The P4–P4′ backbone was guided toward the experimentally observed substrate geometry in the AprX catalytic cleft during energy minimisation.

Each remodelled protein was then positioned against AprX and evaluated for four local criteria: target-backbone geometry, catalytic Zn²⁺ alignment, substrate direction and chain continuity. Whole-protein heavy-atom contacts below 2.6 Å were also recorded as an exploratory steric descriptor. The original analysis treated fewer than 100 contacts as a binary accommodation criterion and used the minimum count across accessible structural states.

To test whether this steric criterion transferred outside AprX, the complete protocol, criteria and thresholds were applied without modification to a thermolysin benchmark comprising 13 native-state test rows from six proteins: seven thermolysin-cleaved rows (six distinct loci because one ambiguous locus was represented by two adjacent assignments), two native-state cross-protease cleavage controls and four resistant rows. Benchmark labels were taken from primary limited-proteolysis studies of the F1-ATPase alpha subunit (Tozawa et al., 1993), D-amino acid oxidase (Campaner et al., 1998),

RNase A (Polverino de Laureto et al., 1997), lysozyme (Polverino de Laureto et al., 1995), and myoglobin (Fontana et al., 1997; Picotti et al., 2004), together with the curated CRBP-II and cross-protease RNase A sites in Hubbard et al. (1998). As an independent sequence check, all benchmark bonds with a readable P1-prime residue matched the hydrophobic thermolysin preference summarised by Hubbard et al. (1998). After the original steric criterion failed this benchmark, contact profiling identified two confounds: contacts from atoms within the deliberately movable window and inflation of raw counts with substrate size. The measure was therefore reformulated as the fraction of substrate atoms outside the movable window that contacted the protease, with the contact distance and remodelling protocol unchanged. This corrected descriptor was then applied to all 33 whey counterexamples and to the nine fully remodelled whey sites.

### 2.15. Statistical analysis and computational packages

Statistical analyses were performed in Python 3.11.16. ROC curves and AUC values were used to test separation between predefined groups. Spearman correlation was used for monotonic associations, Mann–Whitney U tests for continuous two-group comparisons, and Fisher’s exact test for differences in geometry-compatible peptide-pose frequencies. Permutation and bootstrap analyses were used to test robustness; the sequence analysis used 20,000 permutations and 4,000 bootstrap replicates. Conformational-accommodation comparisons also used rank-biserial effect sizes and bootstrap subsampling to account for unequal numbers of casein and whey conformations. For the thermolysin benchmark, ROC AUC and exact label permutation were used to compare cleaved and resistant sites. The corrected distal-contact measure was selected while benchmark labels were visible, so its benchmark performance was treated as development performance rather than independent validation. SciPy was used for statistical testing, scikit-learn for ROC/AUC analysis, NumPy and pandas for numerical calculations and data handling, Biopython/Gemmi for sequence and structural-file handling, Biotite for solvent-accessibility calculations, PDBFixer for protein-structure preparation, and OpenMM for restrained structural minimisation. Additional structural calculations used RDKit, with analyses run locally on an eight-core Windows workstation without GPU or remote computing.

## 3. Results

### 3.1. Structural modelling supports an extended S4 to S4’ substrate footprint for AprX

The AlphaFold AprX model closely matched seven experimentally determined serralysin-family structures. Core Cα RMSD values ranged from 0.60 to 0.73 Å, with sequence identities of 59.5–61.4%. The catalytic region showed high model confidence (mean pLDDT 98.7), and the transferred catalytic Zn²⁺ positions showed a spread of 0.614 Å after structural superposition. Alignment of the substrate-bound 3VI1 structure placed the experimental peptide within the AprX catalytic cleft and defined the central substrate trajectory. Combining 3VI1 with additional serralysin-family ligand structures produced a working S4–S4′ substrate footprint, with strongest structural support for the central positions and weaker support for S3′–S4′ (Figure 3).

**Figure 3.**
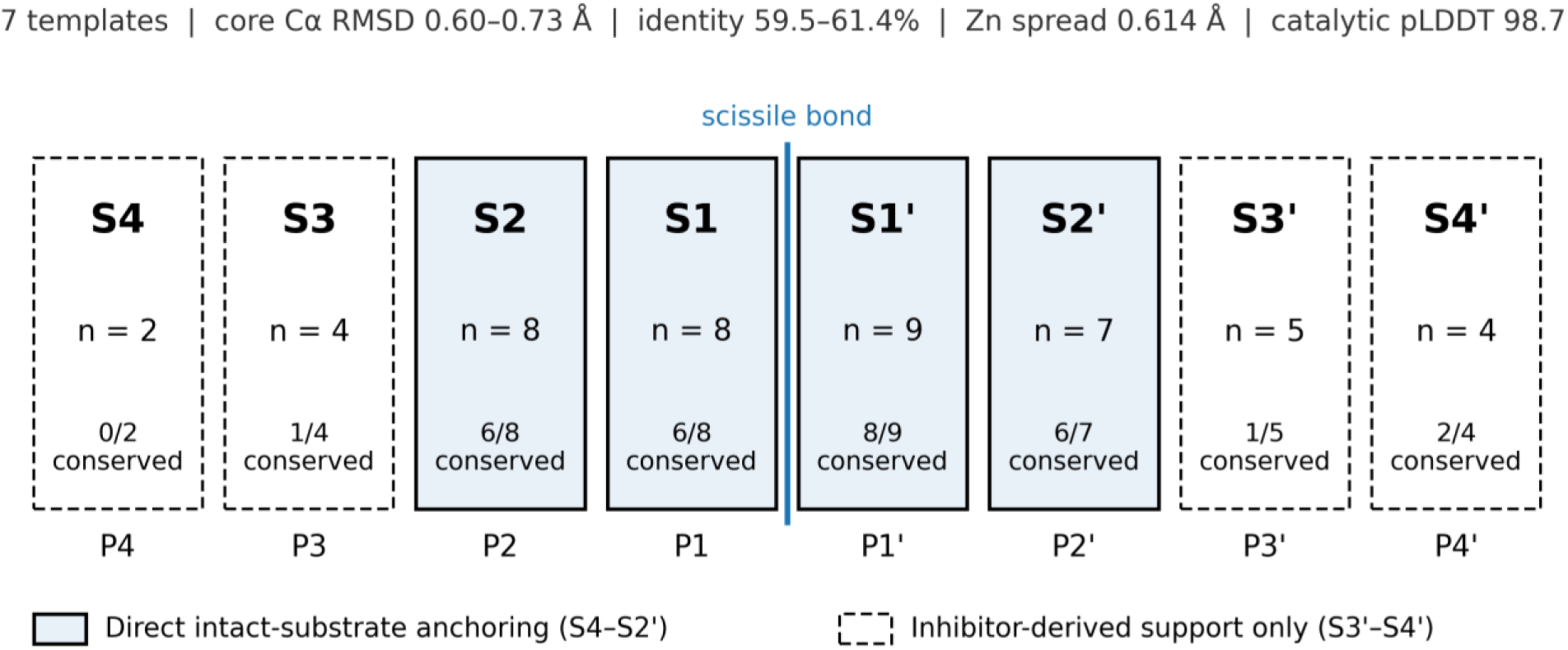
Structural validation and working AprX substrate footprint. The predicted AprX catalytic region was benchmarked against seven serralysin-family structures. Core Cα RMSD, template identity, transferred-Zn consensus and catalytic-region pLDDT are summarised above an S4-S4’ subsite schematic. S4-S2’ are directly anchored by an intact substrate structure, whereas S3’-S4’ are supported by inhibitor-derived primed-side contacts.

### 3.2. Sequence compatibility did not distinguish casein from whey-protein candidate bonds

The dataset produced1,063 candidate peptide bonds, including 780 from caseins and 283 from whey proteins. Sequence-compatibility scores showed little separation between the two protein classes (ROC AUC = 0.522; permutation p = 0.274). β-Lactoglobulin had 14.3% of candidate sites in the highest compatibility tier, compared with 8.9–10.6% across the four caseins. The compatibility score also reproduced several known AprX residue preferences, although discrimination of experimentally observed cleavage sites remained limited (Figure 4).

**Figure 4.**
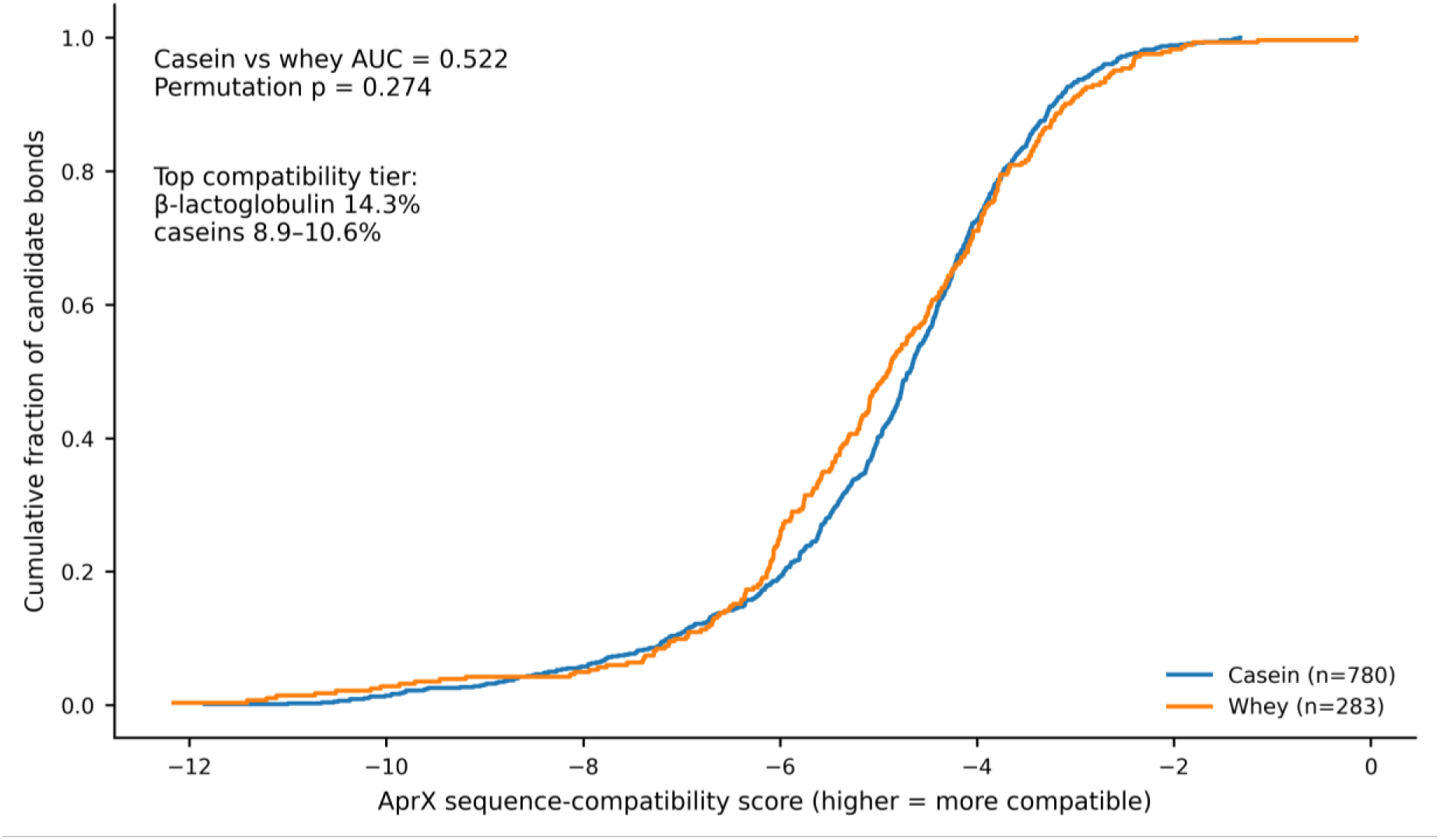
Sequence compatibility does not separate casein from whey candidate bonds. Empirical cumulative distributions of AprX sequence-compatibility scores across all 1,063 candidate bonds. The casein-versus-whey classifier yielded AUC = 0.522 with permutation p = 0.274. β-Lactoglobulin contained 14.3% of candidate bonds in the highest compatibility tier, compared with 8.9-10.6% across caseins.

### 3.3. Accessibility of the immediate cleavage region did not predict AprX cleavage

Local solvent exposure of the scissile region did not distinguish experimentally cleaved casein sites from other candidate sites (AUC = 0.395; permutation p = 0.46). In the whey proteins, 33 candidate sites combined favourable sequence-compatibility scores with local exposure of the scissile bond. These sites were therefore retained for further analysis of accessibility across the complete P4–P4′ substrate window (Figure 5).

**Figure 5.**
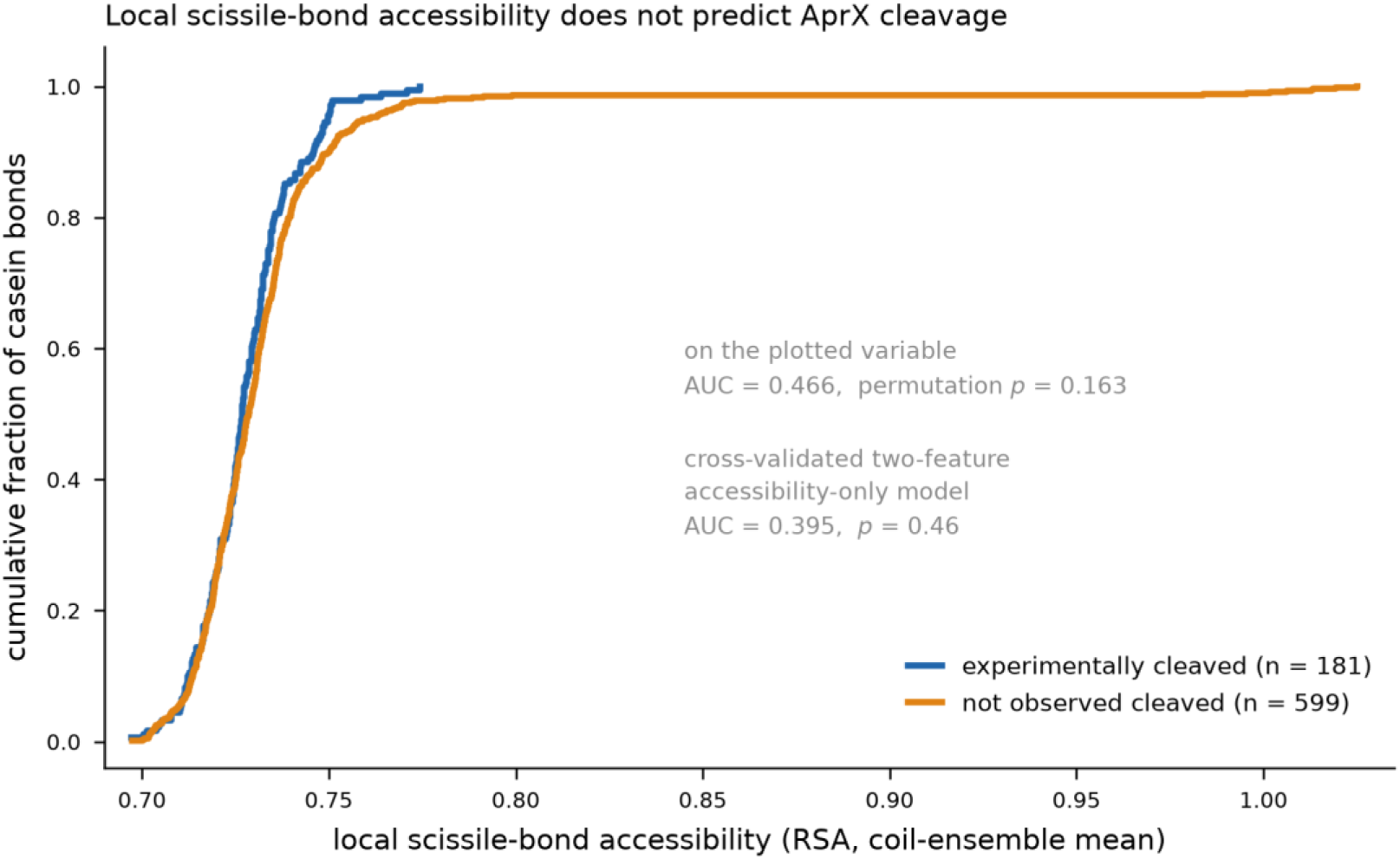
Local scissile-bond accessibility distributions for experimentally cleaved and non-cleaved candidate bonds across the four caseins. The curves show the empirical cumulative distributions of local RSA; inferential statistics for the accessibility-only classifier are reported in the text.

### 3.4. Whey accessibility decreased sharply as the required contiguous substrate window increased

Accessibility decreased sharply as the required contiguous substrate window increased. At an RSA threshold of 0.25, 67 of 281 whey sites were accessible across two residues (23.8%), 15 of 277 across four residues (5.4%), 3 of 273 across six residues (1.1%), and none of 269 across the full eight-residue P4–P4′ window. In contrast, all 756 evaluable casein windows remained accessible at the eight-residue level in the isolated-chain ensemble model. The same overall pattern was observed across all seven RSA thresholds tested (Figure 6).

**Figure 6.**
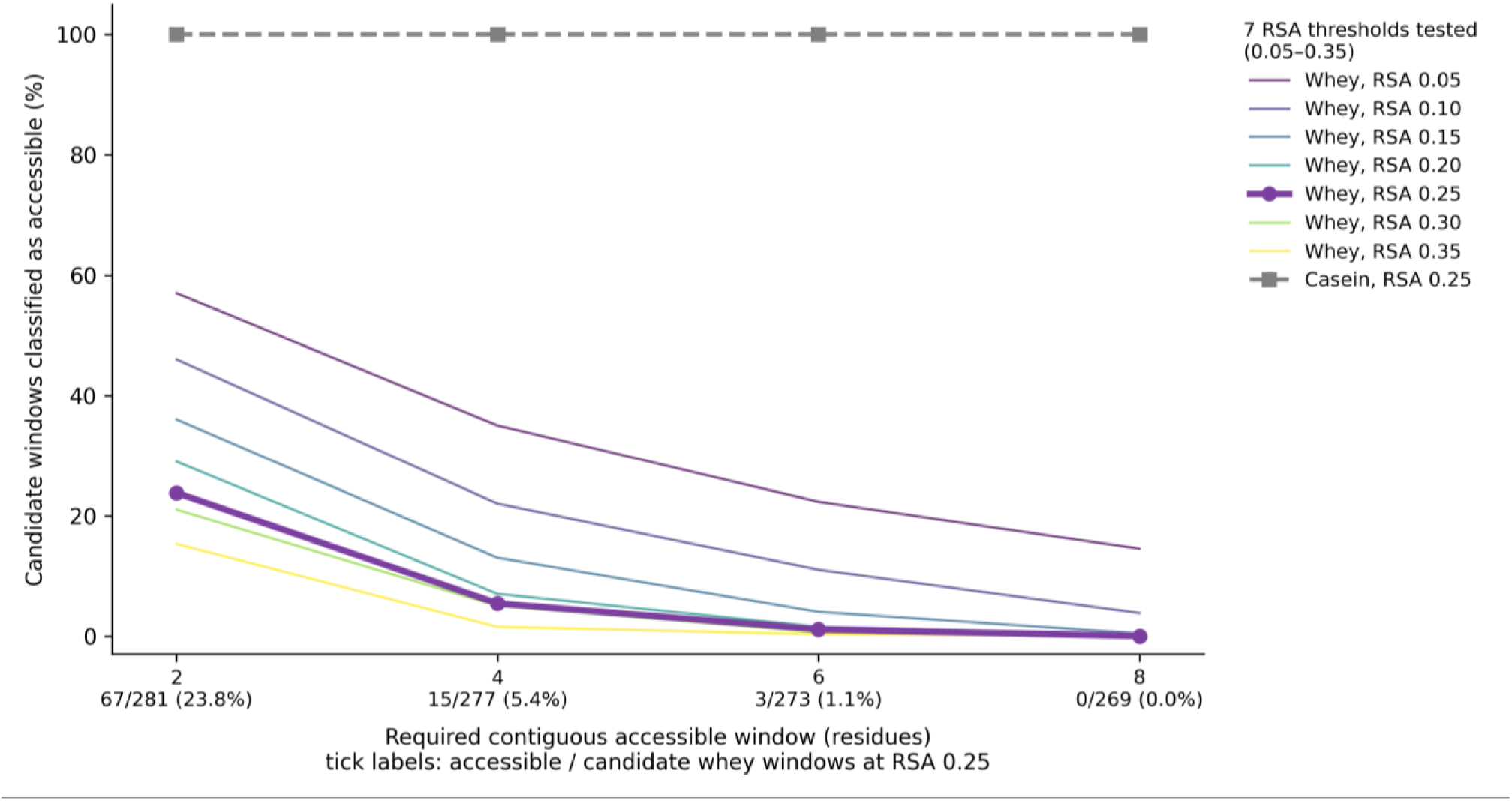
Whey accessibility decreases sharply with increasing contiguous-window length. Percentage of candidate windows classified as accessible across two-, four-, six– and eight-residue requirements. Thin lines show all seven RSA thresholds tested; the RSA = 0.25 series is emphasised. At RSA = 0.25, whey availability decreased from 67/281 (23.8%) to 15/277 (5.4%), 3/273 (1.1%) and 0/269 (0%), while the isolated-chain casein ensemble remained fully accessible.

### 3.5. All 33 sequence-compatible and locally exposed whey sites failed when the full substrate footprint was considered

All 33 whey-protein sites that were both sequence-compatible and locally exposed were re-examined across the full P4–P4′ window. Although the scissile bond was accessible at each site, every window contained at least one neighbouring residue below the accessibility threshold. Consequently, none of the 33 sites remained accessible across the complete eight-residue working footprint (Figure 7).

**Figure 7.**
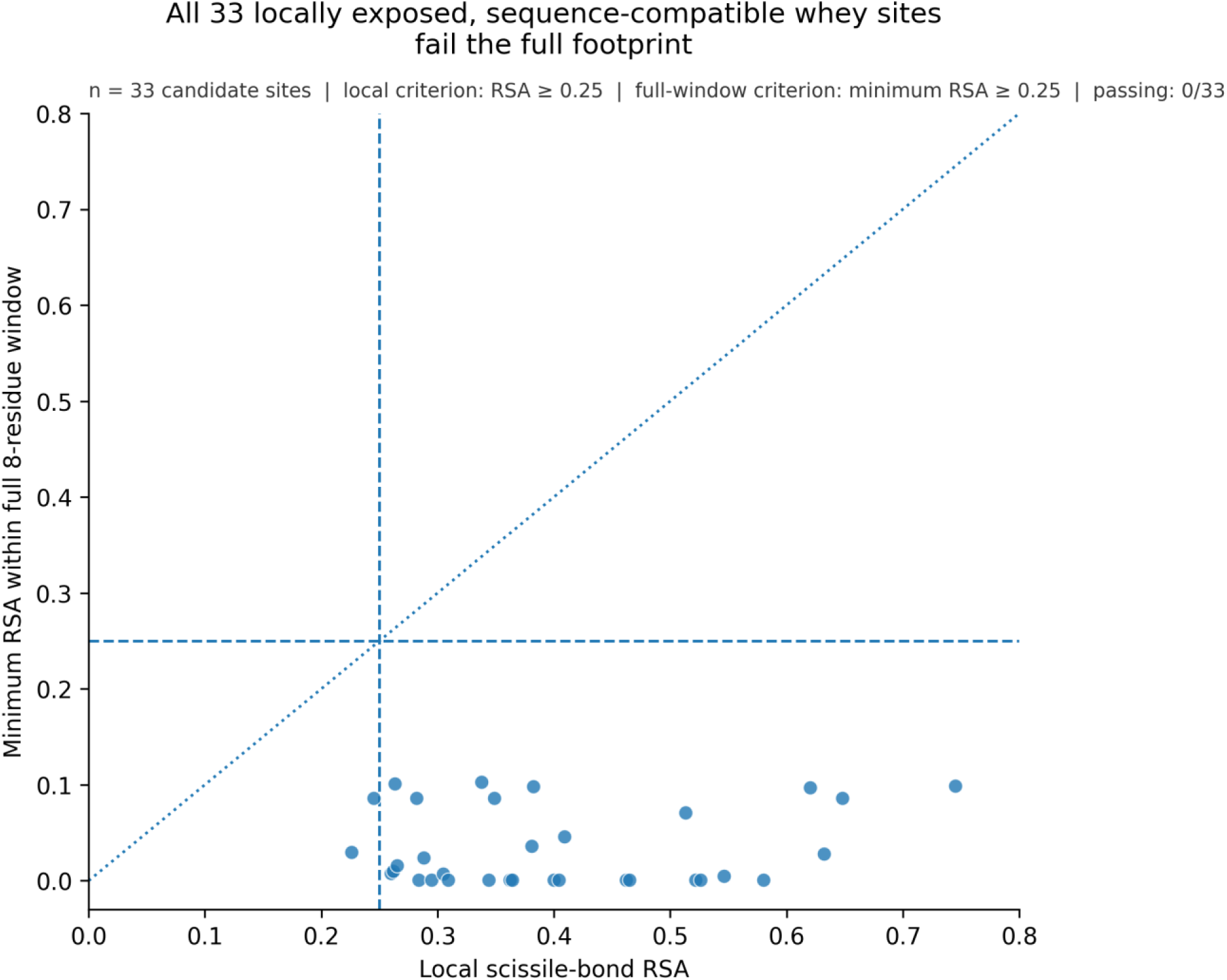
Sequence-compatible and locally exposed whey counterexamples fail at the full substrate footprint. For 33 whey sites meeting both favourable-sequence and local-accessibility criteria, local scissile-bond RSA is plotted against the minimum RSA across the complete eight-residue window. All sites fall below the full-window minimum-RSA criterion of 0.25.

### 3.6. Whey-derived peptide sequences could adopt AprX-compatible catalytic geometries outside the intact protein fold

Peptide threading produced geometry-compatible poses for 5 of 9 whey-derived peptides and 18 of 29 experimentally cleaved casein-derived peptides under the study’s geometric criteria, with no difference between groups by Fisher’s exact test (p = 1.000). The independent 1KAP control reproduced the experimental peptide backbone with an RMSD of approximately 0.73 Å after minimisation, whereas the reversed-orientation control failed. Using the independently supported Zn–O sensitivity limit of 1.84 Å increased geometry-compatible poses to 8 of 9 whey peptides and 25 of 29 casein peptides (Figure 8). These results describe modelled geometry and do not establish catalytic turnover.

**Figure 8.**
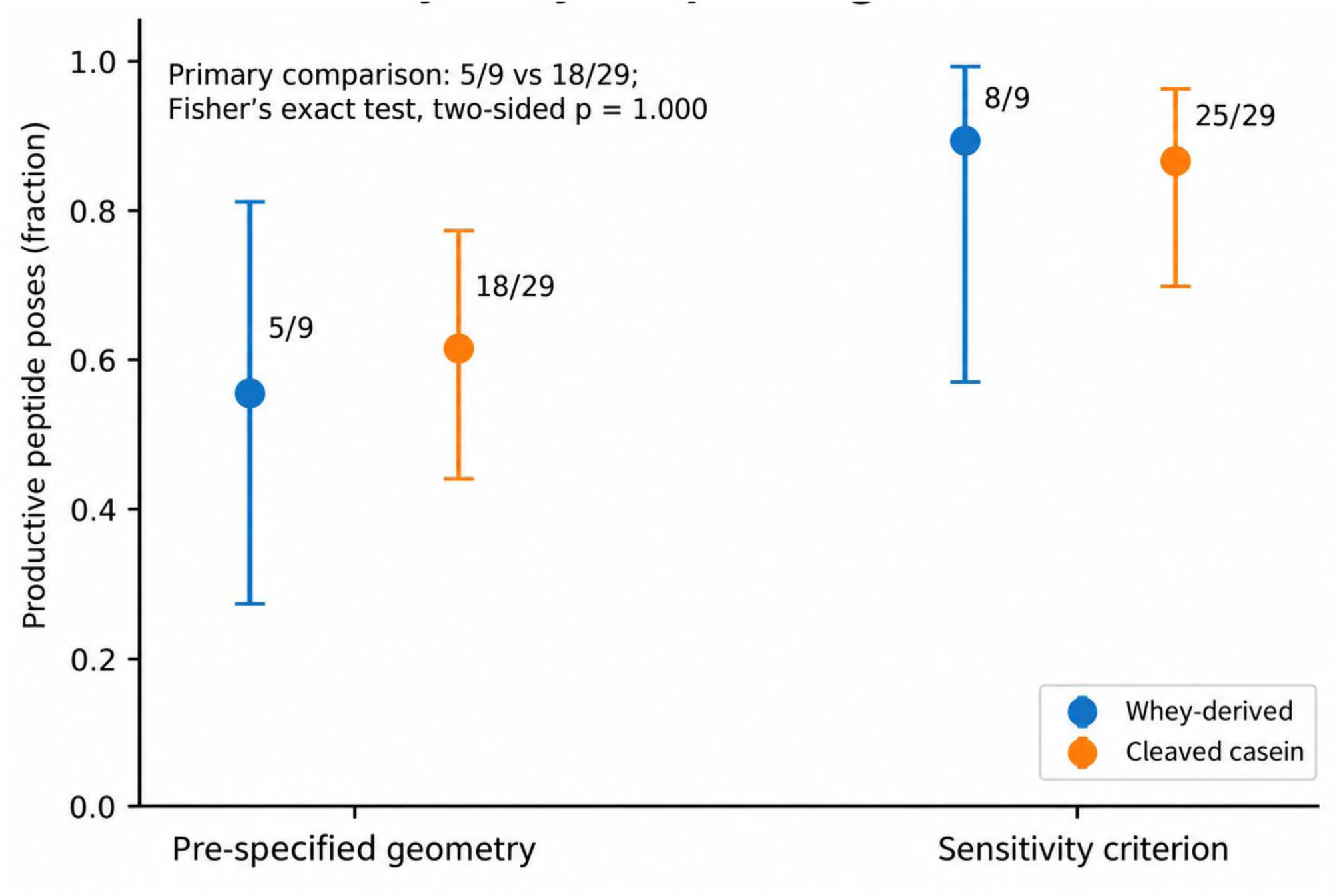
Whey-derived peptides can adopt AprX-compatible modelled geometries outside the intact fold. Geometry-compatible pose fractions are shown for whey-derived peptides and experimentally cleaved casein peptides using the original pre-specified criteria and the independently justified sensitivity criterion. Error bars are Wilson 95% confidence intervals. The primary comparison was 5/9 versus 18/29, Fisher’s exact two-sided p = 1.000.

### 3.7. Local backbone accommodation was retained, but external benchmarking rejected rigid-placement steric exclusion

Conformational-accommodation analysis showed similar minimum rearrangement requirements for casein and whey sites, with a median minimum accommodation window of 12 residues in both groups (p = 0.058). With larger movable windows, whey segments approached the target AprX-bound geometry, reaching a median best backbone RMSD of 0.79 Å while maintaining chain closure and correct substrate direction. The P1 carbonyl oxygen was also positioned within catalytic distance of Zn²⁺ (median 2.14 Å) (Figure 9). The local part of the protocol was also supported by the external thermolysin benchmark, in which 12 of 13 rows satisfied all four local criteria.

**Figure 9.**
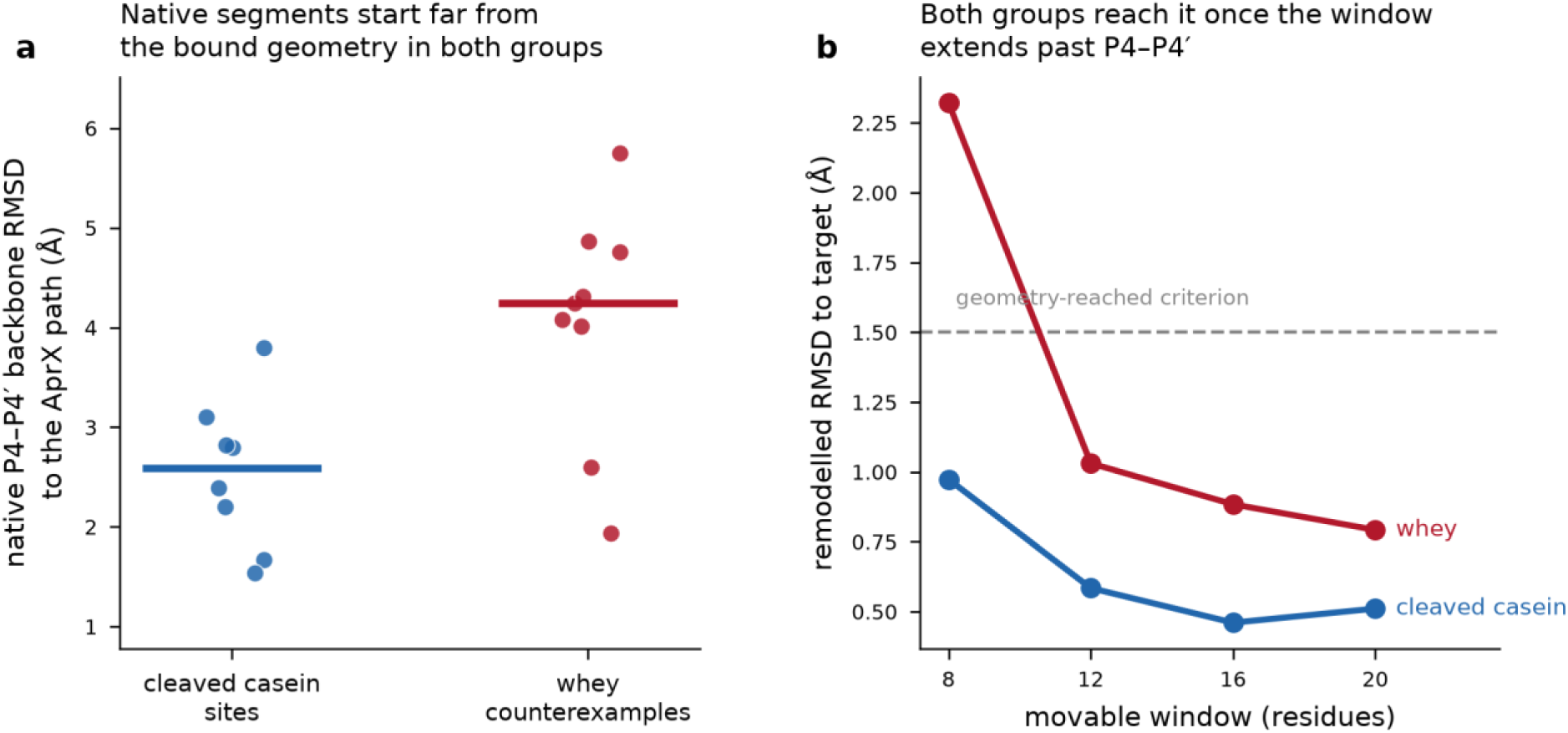
Local backbone accommodation of experimentally cleaved casein sites and whey counterexamples. (a) Native P4–P4′ segments in both groups initially deviate from the AprX-bound substrate path. (b) Both groups approach the target geometry as the movable window increases.

By contrast, the original whole-protein steric criterion failed external benchmarking. None of the 13 thermolysin benchmark rows met the <100-contact criterion, including 0 of 7 rows known to be cleaved by thermolysin (Tozawa et al., 1993; Hubbard et al., 1998; Campaner et al., 1998) and 0 of 2 cross-protease controls (Hubbard et al., 1998), although 12 of 13 passed the four local criteria. Raw contact count poorly discriminated thermolysin-cleaved from resistant sites (AUC = 0.571), and a threshold sweep did not identify a significant alternative cutoff.

Contact profiling showed that the original score penalised atoms within the remodelled movable window and increased with substrate size. After excluding movable-window atoms and expressing contacts as the fraction of distal substrate atoms in contact, the seven thermolysin-cleaved rows spanned 0.024–0.445 and the four resistant rows 0.574–0.766 (AUC = 1.000; exact permutation p = 0.003). Because this corrected measure was selected with the benchmark labels visible, these values represent development performance, not independent validation; including the two cross-protease controls reduced the AUC to 0.889.

When the corrected measure was applied back to the AprX whey sites, 22 of 33 native placements fell within the thermolysin-cleaved range, six fell in the gap and five in the resistant range. Among the nine fully remodelled whey sites, five fell in the cleaved range, three in the gap and one in the resistant range. Median values for the 33 native and nine remodelled whey sets were 0.331 and 0.337, respectively, and no β-lactoglobulin site reached the resistant range. Remodelling improved P4–P4′ fit from a median 4.25 Å to 1.04 Å while changing the corrected steric measure little (r = 0.988; maximum change 0.063). The corrected analysis therefore did not restore the original whole-protein steric-exclusion claim (Figures 10–12).

**Figure 10.**
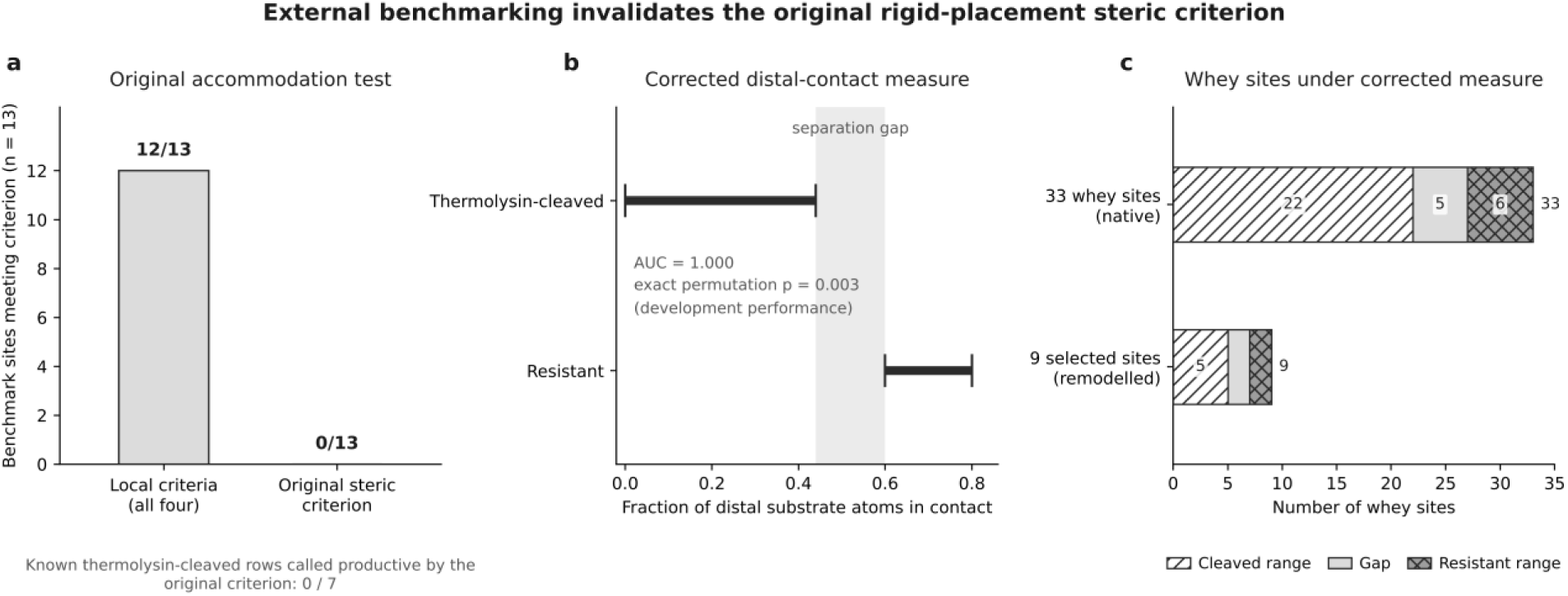
External benchmark and restoration test for the whole-protein steric descriptor. (a) In the thermolysin benchmark, 12 of 13 rows passed the four local criteria, whereas none passed the original steric criterion and 0 of 7 thermolysin-cleaved rows were called productive. (b) After excluding the movable window and normalising contacts by distal substrate size, thermolysin-cleaved and resistant sites separated in the development set (AUC = 1.000; exact permutation p = 0.003). (c) Applying the corrected measure to AprX whey sites placed most native and remodelled sites within the thermolysin-cleaved range or the intervening gap rather than the resistant range.

**Figure 11.**
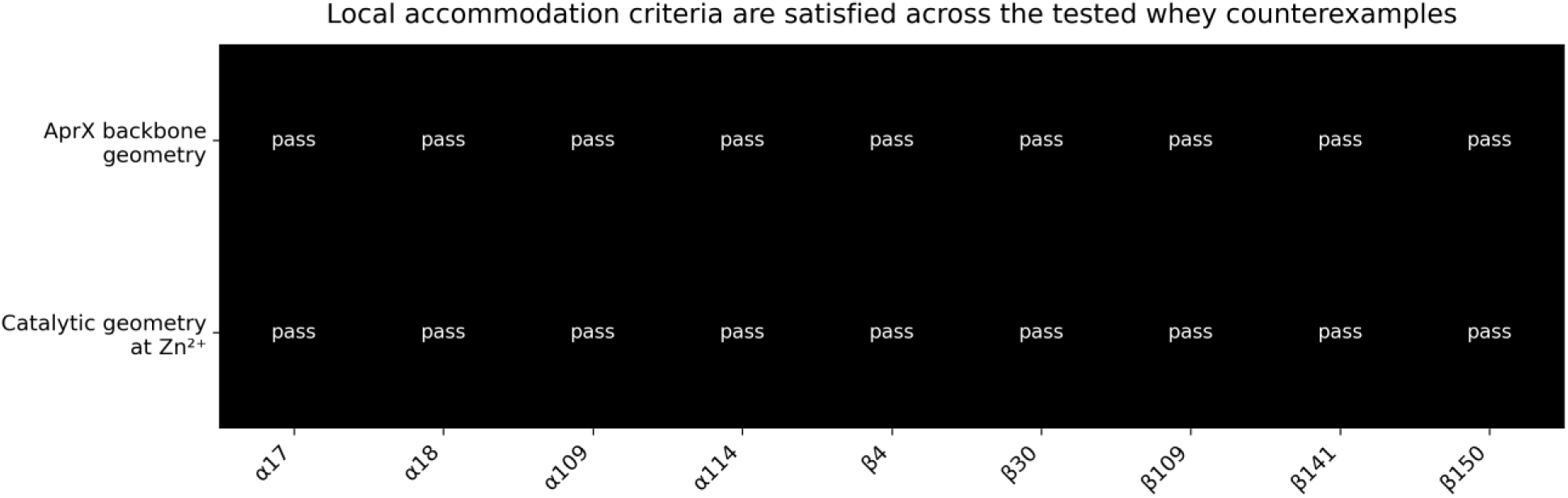
Local accommodation criteria for the nine tested whey counterexamples. Each site reached the AprX substrate-backbone geometry and catalytic Zn²⁺ geometry after remodelling. The original whole-protein steric pass/fail row was removed because external benchmarking showed that the absolute steric criterion was over-conservative.

**Figure 12.**
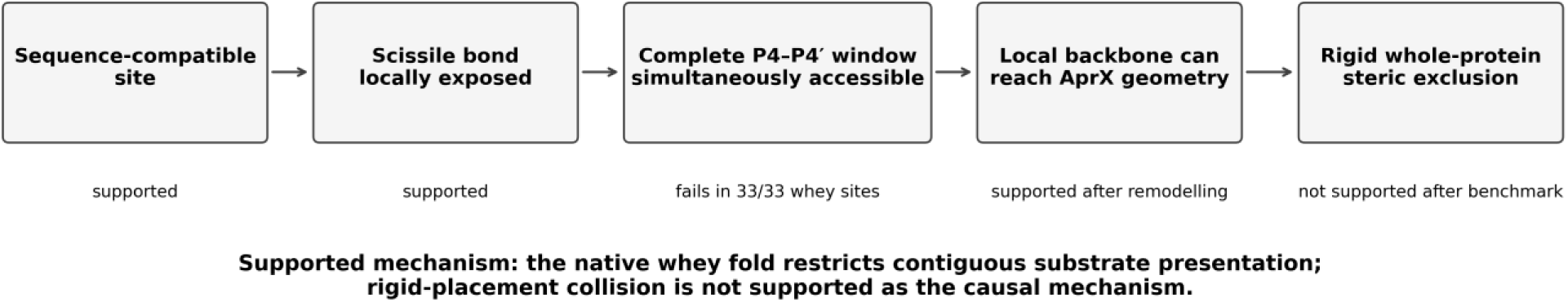
Revised summary after external benchmarking. Sequence compatibility and local scissile-bond exposure did not separate the protein classes, and all 33 whey counterexamples failed the complete P4–P4′ accessibility requirement. Local remodelling showed that the selected whey segments could attain AprX-compatible geometry. External benchmarking did not support the original rigid whole-protein steric interpretation. The remaining structural result is the loss of contiguous substrate-window accessibility in the native whey structures used here.

## 4. Discussion

Native whey proteins are less susceptible to AprX than caseins. This difference has usually been linked to the compact whey fold, while caseins are intrinsically disordered (Matéos et al., 2015; Holt et al., 2013). The present study defines this structural difference more precisely. Sequence composition, scissile-bond exposure and local backbone flexibility did not separate the two protein classes. The clearest difference appeared when accessibility was measured across a contiguous substrate-sized region. At the principal RSA threshold, whey accessibility fell from 23.8% over two residues to 0% across the P4–P4′ working footprint. All evaluable casein windows remained accessible in the isolated-chain ensembles. The same trend remained across all tested RSA thresholds. The main result is therefore that native whey folds restrict simultaneous exposure across the region that AprX is expected to engage.

This result is important because AprX has broad sequence tolerance. Matéos et al. (2015) mapped many cleavage sites across αS1-, β– and κ-caseins. Zhang et al. (2023) also found position-dependent preferences without a strict cleavage motif. Our analysis tested whether favourable sites were less common in whey proteins. They were not. The casein-versus-whey classifier gave an AUC of 0.522. β-Lactoglobulin also contained 14.3% of candidate sites in the highest compatibility tier, compared with 8.9–10.6% in the individual caseins. Sequence may therefore help determine which bond AprX selects once a region is available, but it does not explain the large difference in whole-protein susceptibility between caseins and whey proteins. This agrees with the broader prediction literature. Barkan et al. (2010) found that combining sequence and structural features improved prediction of protease substrates over sequence-only models. Their work treated structural context as an added predictor of cleavage. The present data support that separation for AprX, but identify a more specific structural variable at the scale of the substrate footprint.

Local solvent exposure gave a similar result. Kazanov et al. (2011) and Belushkin et al. (2014) showed that exposed regions are enriched among proteolytic sites across many proteases. Novotný and Bruccoleri (1987) went further and calculated accessibility using probes comparable in size to proteolytic enzymes. They found that known proteolytic sites were highly accessible to these larger probes, while flexibility was not required at every site. In the present dataset, local RSA did not distinguish observed casein cleavage sites from other candidate bonds. In addition, 33 whey sites were both sequence-compatible and locally exposed. These sites show that residue-level RSA is not enough in this system. A bond can be exposed to solvent while neighbouring residues remain unavailable as one contiguous substrate segment.

The window analysis added information that was not captured by residue-level accessibility. As more neighbouring residues were included, accessibility in whey proteins fell from 23.8% to 5.4%, 1.1% and finally 0%. This pattern agrees with the extended binding mode seen in protease structures (Madala et al., 2010). It also agrees with the strong effect of peptide length on serralysin hydrolysis reported by Louis et al. (1998). The new point is not that serralysin uses several subsites. That is already known. The new point is that this requirement reveals a structural constraint in intact whey proteins that is almost invisible when only the cleavage bond is examined.

The local-remodelling analysis narrowed the mechanism further. Hubbard et al. (1994) showed that cleavage of folded proteins can require local backbone rearrangement before a substrate can adopt a protease-compatible geometry. Park and Marqusee (2004) reached a related conclusion using native-state proteolysis. They showed that cleavage can occur from transient higher-energy conformations that expose a normally protected region without global unfolding. These studies make local conformational access a plausible explanation for whey resistance. Our data did not support that explanation as the main difference between casein and whey. Both groups had a median minimum accommodation window of 12 residues. Most remodelled whey segments also reached the target backbone and catalytic Zn²⁺ geometry. The local whey segment was therefore not uniquely unable to adopt the required conformation. The stronger constraint appeared before this step, in the simultaneous accessibility of the contiguous region within the native fold.

The external thermolysin benchmark tested a stronger interpretation of this result. The original rigid-contact metric gave a large casein-whey separation. However, it also rejected all seven benchmark sites known to be cleaved by thermolysin. Contact profiling showed two clear problems. The metric counted atoms from the remodelled region itself. It was also affected by substrate size. After correction, the distal-contact measure separated cleaved and resistant thermolysin sites in the development set. However, it did not classify most AprX whey sites as resistant. The benchmark therefore removed support for a rigid whole-protein collision mechanism. It did not change the sequence result, the local-RSA result, the contiguous-window collapse, the peptide-threading result or the local-remodelling result. The supported conclusion is narrower. Native whey structures restrict simultaneous presentation of the working substrate footprint, but the present analysis does not show that whole-protein steric collision prevents cleavage.

This distinction matters for current proteolysis prediction. Matveev et al. (2023) developed a structure-based predictor that uses solvent accessibility, secondary structure, B-factor, loop length and terminal flexibility. These features describe the local structural environment of each peptide bond. Haack and Kalogeropoulos (2025) also recommend SASA as a structural measure in positional proteomics and describe it as a proxy for whether a cleavage site is surface exposed. These approaches are useful, but they still evaluate structural susceptibility mainly at or near the candidate bond. The AprX results show that an additional signal can appear only when several neighbouring residues are required to be accessible at the same time.

Recent work also shows why this distinction may matter for prediction accuracy. Yu et al. (2026) combined experimentally derived protease sequence preferences with AlphaFold-based structural features in PhageScout. The structural variables included accessibility and flexibility around candidate sites. Adding these features gave only modest and protease-dependent improvements over sequence-derived information. The present study does not explain that result, but it identifies one structural feature that such site-centred descriptors do not measure directly. Footprint-scale accessibility asks whether the complete region needed for enzyme engagement is simultaneously available, not only whether the central bond or nearby residues appear exposed.

The result also changes how structural susceptibility can be framed. Current models often assign a score to each peptide bond from local sequence and local structural features. Barkan et al. (2010), Matveev et al. (2023), Haack and Kalogeropoulos (2025), and Yu et al. (2026) represent increasingly structure-aware versions of this approach. The present data suggest that a second structural scale can be useful. One score can describe the candidate bond. A separate score can describe simultaneous accessibility across the expected protease-binding footprint. These two quantities should not be assumed to carry the same information.

This framework may also matter when processing changes tertiary structure without changing sequence. Heating, ligand loss or partial unfolding could increase access across a candidate substrate region. Aggregation could reduce it. Guo et al. (1995) showed that structural perturbation increased β-lactoglobulin susceptibility to other proteases. For AprX, the direct prediction is narrower. Treatments that increase simultaneous exposure across a candidate P4–P4′ region should increase access to that region. Whether this also increases cleavage must be tested experimentally.

The casein ensembles represent isolated disordered chains. They do not represent casein micelles with intermolecular contacts and colloidal calcium phosphate (Holt et al., 2013). Therefore, 756 of 756 accessible casein windows does not mean that every casein segment is exposed in milk. It means only that the isolated casein chains did not show the persistent tertiary occlusion seen in the experimental whey structures. The P4–P4′ interval is also a working structural footprint and should not be interpreted as a proven minimum substrate length.

To summarise, single-residue RSA should not be treated as equivalent to substrate accessibility in this system. Accessibility should also be measured across the region expected to occupy the protease cleft. This extends, rather than replaces, current sequence– and structure-based prediction approaches. Rigid whole-protein clash scores should also not be used as cleavage filters without external positive and negative controls. The thermolysin benchmark showed that a metric can separate two groups within one dataset and still fail on known cleavage sites. The result that remains after that test is narrower but stronger. AprX resistance in native whey proteins is associated with failure to present a contiguous substrate-sized region, rather than with a shortage of favourable sequences, burial of the scissile bond alone, or an inability of the local peptide backbone to reach the bound geometry.

### 4.1. Implications, limitations and experimental validation

The main limitations are tied to the claims above. AprX was represented by an AlphaFold model, although its catalytic region agreed closely with experimental serralysin structures. P3′–P4′ had weaker structural support than the central subsites, so the P4–P4′ interval is a working footprint rather than a fixed minimum. The contiguous-window analysis was post-hoc. The casein ensembles did not model micellar organisation. The corrected distal-contact descriptor was developed with thermolysin labels visible and failed one cross-protease control, so it is not independently validated. The study also did not measure turnover at the modelled whey sites. A direct test would compare matched native and structurally perturbed whey proteins under the same AprX conditions and map cleavage at the predicted counterexample sites. If full-window exposure increased without a corresponding increase in cleavage, the contiguous-accessibility interpretation would weaken. Flexible whole-protein encounter effects remain unresolved.

## Data availability

All data supporting the findings of this study are provided within the article and its Supplementary Tables. The Supplementary Tables include the datasets used for sequence scoring, structural accessibility, substrate-window analysis, peptide modelling, conformational remodelling and external thermolysin benchmarking.

## References

1. Almeida, L. F., Machado, S. G., Ventorim, R. Z., de Carvalho, A. F., and Maitan-Alfenas, G. P. 2026. Unveiling AprX: biochemical and molecular mechanisms of a heat-resistant inducing milk spoilage protease. Food Research International, 230, Article 118553. 10.1016/j.foodres.2026.118553.

2. Baglinière, F., Matéos, A., Tanguy, G., Jardin, J., Briard-Bion, V., Rousseau, F., Robert, B., Beaucher, E., Gaillard, J. L., Amiel, C., Humbert, G., Dary, A., and Gaucheron, F. 2013. Proteolysis of ultra high temperature-treated casein micelles by AprX enzyme from *Pseudomonas fluorescens* F induces their destabilisation. International Dairy Journal, 31, 55–61. 10.1016/j.idairyj.2013.02.011.

3. Barkan, D. T., Hostetter, D. R., Mahrus, S., Pieper, U., Wells, J. A., Craik, C. S., and Sali, A. 2010. Prediction of protease substrates using sequence and structure features. Bioinformatics, 26, 1714–1722. 10.1093/bioinformatics/btq267.

4. Belushkin, A. A., Vinogradov, D. V., Gelfand, M. S., Osterman, A. L., Cieplak, P., and Kazanov, M. D. 2014. Sequence-derived structural features driving proteolytic processing. Proteomics, 14, 42–50. 10.1002/pmic.201300416.

5. Campaner, S., Pollegioni, L., Ross, B. D., and Pilone, M. S. 1998. Limited proteolysis and site-directed mutagenesis reveal the origin of microheterogeneity in *Rhodotorula gracilis* D-amino acid oxidase. Biochemical Journal, 330, 615–621. 10.1042/bj3300615.

6. Eastman, P., Swails, J., Chodera, J. D., McGibbon, R. T., Zhao, Y., Beauchamp, K. A., Wang, L.-P., Simmonett, A. C., Harrigan, M. P., Stern, C. D., Wiewiora, R. P., Brooks, B. R., and Pande, V. S. 2017. OpenMM 7: rapid development of high performance algorithms for molecular dynamics. PLOS Computational Biology, 13, Article e1005659. 10.1371/journal.pcbi.1005659.

7. Ellison, D., Hinton, J., Hubbard, S. J., and Beynon, R. J. 1995. Limited proteolysis of native proteins: the interaction between avidin and proteinase K. Protein Science, 4, 1337–1345. 10.1002/pro.5560040709.

8. Fontana, A., Zambonin, M., Polverino de Laureto, P., De Filippis, V., Clementi, A., and Scaramella, E. 1997. Probing the conformational state of apomyoglobin by limited proteolysis. Journal of Molecular Biology, 266, 223–230. 10.1006/jmbi.1996.0787.

9. Fontana, A., Polverino de Laureto, P., Spolaore, B., Frare, E., Picotti, P., and Zambonin, M. 2004. Probing protein structure by limited proteolysis. Acta Biochimica Polonica, 51, 299–321. 10.18388/abp.2004_3573.

10. Guo, M. R., Fox, P. F., Flynn, A., and Kindstedt, P. S. 1995. Susceptibility of beta-lactoglobulin and sodium caseinate to proteolysis by pepsin and trypsin. Journal of Dairy Science, 78, 2336–2344. 10.3168/jds.S0022-0302(95)76860-6.

11. Haack, A. M., and Kalogeropoulos, K. 2025. Data processing and analysis in positional proteomics. Proteomics, 25, 277–293. 10.1002/pmic.70069.

12. Holt, C., Carver, J. A., Ecroyd, H., and Thorn, D. C. 2013. Invited review: caseins and the casein micelle: their biological functions, structures, and behavior in foods. Journal of Dairy Science, 96, 6127–6146. 10.3168/jds.2013-6831.

13. Hubbard, S. J., Eisenmenger, F., and Thornton, J. M. 1994. Modeling studies of the change in conformation required for cleavage of limited proteolytic sites. Protein Science, 3, 757–768. 10.1002/pro.5560030505.

14. Hubbard, S. J., Beynon, R. J., and Thornton, J. M. 1998. Assessment of conformational parameters as predictors of limited proteolytic sites in native protein structures. Protein Engineering, 11, 349–359. 10.1093/protein/11.5.349.

15. Huey, R., Morris, G. M., Olson, A. J., and Goodsell, D. S. 2007. A semiempirical free energy force field with charge-based desolvation. Journal of Computational Chemistry, 28, 1145–1152. 10.1002/jcc.20634.

16. Jumper, J., Evans, R., Pritzel, A., et al. 2021. Highly accurate protein structure prediction with AlphaFold. Nature, 596, 583–589. 10.1038/s41586-021-03819-2.

17. Kabsch, W. 1976. A solution for the best rotation to relate two sets of vectors. Acta Crystallographica Section A, 32, 922–923. 10.1107/S0567739476001873.

18. Kazanov, M. D., Igarashi, Y., Eroshkin, A. M., Cieplak, P., Ratnikov, B., Zhang, Y., Li, Z., Godzik, A., Osterman, A. L., and Smith, J. W. 2011. Structural determinants of limited proteolysis. Journal of Proteome Research, 10, 3642–3651. 10.1021/pr200271w.

19. Kontopidis, G., Holt, C., and Sawyer, L. 2004. Invited review: beta-lactoglobulin: binding properties, structure, and function. Journal of Dairy Science, 87, 785–796. 10.3168/jds.S0022-0302(04)73222-1.

20. Kunzmann, P., and Hamacher, K. 2018. Biotite: a unifying open source computational biology framework in Python. BMC Bioinformatics, 19, Article 346. 10.1186/s12859-018-2367-z.

21. Louis, D., Bernillon, J., and Wallach, J. M. 1998. Specificity of *Pseudomonas aeruginosa* serralysin revisited, using biologically active peptides as substrates. Biochimica et Biophysica Acta, 1387, 378–386. 10.1016/S0167-4838(98)00144-7.

22. Madala, P. K., Tyndall, J. D. A., Nall, T., and Fairlie, D. P. 2010. Update 1 of: proteases universally recognize beta strands in their active sites. Chemical Reviews, 110, PR1–PR31. 10.1021/cr900368a.

23. Marchand, S., Vandriesche, G., Coorevits, A., Coudijzer, K., De Jonghe, V., Dewettinck, K., De Vos, P., Devreese, B., Heyndrickx, M., and De Block, J. 2009. Heterogeneity of heat-resistant proteases from milk *Pseudomonas* species. International Journal of Food Microbiology, 133, 68–77. 10.1016/j.ijfoodmicro.2009.04.027.

24. Matéos, A., Guyard-Nicodème, M., Baglinière, F., Jardin, J., Gaucheron, F., Dary, A., Humbert, G., and Gaillard, J.-L. 2015. Proteolysis of milk proteins by AprX, an extracellular protease identified in *Pseudomonas* LBSA1 isolated from bulk raw milk, and implications for the stability of UHT milk. International Dairy Journal, 49, 78–88. 10.1016/j.idairyj.2015.04.008.

25. Matveev, E. V., Safronov, V. V., Ponomarev, G. V., and Kazanov, M. D. 2023. Predicting structural susceptibility of proteins to proteolytic processing. International Journal of Molecular Sciences, 24, Article 10761. 10.3390/ijms241310761.

26. Miller, S., Janin, J., Lesk, A. M., and Chothia, C. 1987. Interior and surface of monomeric proteins. Journal of Molecular Biology, 196, 641–656. 10.1016/0022-2836(87)90038-6.

27. Morris, G. M., Huey, R., Lindstrom, W., Sanner, M. F., Belew, R. K., Goodsell, D. S., and Olson, A. J. 2009. AutoDock4 and AutoDockTools4: automated docking with selective receptor flexibility. Journal of Computational Chemistry, 30, 2785–2791. 10.1002/jcc.21256.

28. Novotný, J., and Bruccoleri, R. E. 1987. Correlation among sites of limited proteolysis, enzyme accessibility and segmental mobility. FEBS Letters, 211, 185–189. 10.1016/0014-5793(87)81433-3.

29. Ozenne, V., Bauer, F., Salmon, L., Huang, J.-R., Jensen, M. R., Segard, S., Bernadó, P., Charavay, C., and Blackledge, M. 2012. Flexible-meccano: a tool for the generation of explicit ensemble descriptions of intrinsically disordered proteins and their associated experimental observables. Bioinformatics, 28, 1463–1470. 10.1093/bioinformatics/bts172.

30. Park, C., and Marqusee, S. 2004. Probing the high energy states in proteins by proteolysis. Journal of Molecular Biology, 343, 1467–1476. 10.1016/j.jmb.2004.08.085.

31. Permyakov, E. A., Permyakov, S. E., Breydo, L., Redwan, E. M., Almehdar, H. A., and Uversky, V. N. 2016. Disorder in milk proteins: alpha-lactalbumin. Part A. Structural properties and conformational behavior. Current Protein & Peptide Science, 17, 352–367. 10.2174/138920371704160405001222.

32. Picotti, P., Marabotti, A., Negro, A., Musi, V., Spolaore, B., Zambonin, M., and Fontana, A. 2004. Modulation of the structural integrity of helix F in apomyoglobin by single amino acid replacements. Protein Science, 13, 1572–1585. 10.1110/ps.04635304.

33. Polverino de Laureto, P., De Filippis, V., Scaramella, E., Zambonin, M., and Fontana, A. 1995. Limited proteolysis of lysozyme in trifluoroethanol: isolation and characterization of a partially active enzyme derivative. European Journal of Biochemistry, 230, 779–787. 10.1111/j.1432-1033.1995.0779h.x.

34. Polverino de Laureto, P., Scaramella, E., De Filippis, V., Bruix, M., Rico, M., and Fontana, A. 1997. Limited proteolysis of ribonuclease A with thermolysin in trifluoroethanol. Protein Science, 6, 860–872. 10.1002/pro.5560060413.

35. Qin, B. Y., Bewley, M. C., Creamer, L. K., Baker, E. N., and Jameson, G. B. 1999. Functional implications of structural differences between variants A and B of bovine beta-lactoglobulin. Protein Science, 8, 75–83. 10.1110/ps.8.1.75.

36. Shrake, A., and Rupley, J. A. 1973. Environment and exposure to solvent of protein atoms. Lysozyme and insulin. Journal of Molecular Biology, 79, 351–371. 10.1016/0022-2836(73)90011-9.

37. Tien, M. Z., Meyer, A. G., Sydykova, D. K., Spielman, S. J., and Wilke, C. O. 2013. Maximum allowed solvent accessibilities of residues in proteins. PLOS ONE, 8, Article e80635. 10.1371/journal.pone.0080635.

38. Tozawa, K., Miyauchi, M., and Yoshida, M. 1993. Structure of the alpha subunit of F1-ATPase probed by limited proteolysis. Journal of Biological Chemistry, 268, 19044–19054. 10.1016/S0021-9258(17)46733-0.

39. Varadi, M., Anyango, S., Deshpande, M., et al. 2022. AlphaFold Protein Structure Database: massively expanding the structural coverage of protein-sequence space with high-accuracy models. Nucleic Acids Research, 50, D439–D444. 10.1093/nar/gkab1061.

40. Yu, E., Holding, M. L., Huang, R., Chan, A., Teney, C., and Kretz, C. A. 2026. PhageScout: protease cleavage site prediction using an experimental substrate phage display motif-based approach. International Journal of Molecular Sciences, 27, Article 7593. 10.3390/ijms27177593.

41. Zhang, C., Bijl, E., and Hettinga, K. 2018. Destabilization of UHT milk by protease AprX from *Pseudomonas fluorescens* and plasmin. Food Chemistry, 263, 127–134. 10.1016/j.foodchem.2018.04.128.

42. Zhang, C., Bijl, E., Svensson, B., and Hettinga, K. 2019. The extracellular protease AprX from *Pseudomonas* and its spoilage potential for UHT milk: a review. Comprehensive Reviews in Food Science and Food Safety, 18, 834–852. 10.1111/1541-4337.12452.

43. Zhang, C., Bijl, E., Muis, K. E., and Hettinga, K. 2020. Stability of fat globules in UHT milk during proteolysis by the AprX protease from *Pseudomonas fluorescens* and by plasmin. Journal of Dairy Science, 103, 179–190. 10.3168/jds.2019-17150.

44. Zhang, C., Boeren, S., Zhao, L., Bijl, E., and Hettinga, K. 2023. The impact of low-temperature inactivation of protease AprX from *Pseudomonas* on its proteolytic capacity and specificity: a peptidomic study. Dairy, 4, 150–166. 10.3390/dairy4010011.

